# Functional Assembly of the Qc-SNARE with Sec18 and Sec17 on Membranes

**DOI:** 10.1101/2025.10.21.683730

**Authors:** Amy Orr, Karina Lopes, Jerry O’Dwyer, William Wickner

## Abstract

Yeast vacuolar fusion is driven by Sec17, Sec18, SNAREs (R, Qa, Qb, Qc) and HOPS, a catalyst of SNARE assembly. Qc, the only vacuolar SNARE that is not membrane-anchored, has a unique path of assembly with other fusion catalysts. Qc is the only SNARE which binds Sec17 with high affinity. Sec18 confers a high affinity for Qc (but not Qb) on HOPS-dependent fusion, but it has been unclear how Sec18 acts. The membrane complex of Sec17 and Sec18 bind Qc to form a membrane:Sec18:Sec17:Qc complex. Sec18 ATP hydrolysis, though dispensable for fusion, provides a measure of the physical and functional interactions between Qc, Sec17, Sec18, and membranes. Each binary interface in this quaternary complex regulates Sec18 ATPase and fusion. Qc is better than other SNAREs, alone or in combination, for stimulating ATP hydrolysis. We propose a working model in which membrane-bound Qc:Sec17:Sec18 associates with the *trans* complex of HOPS:R:QaQb, displacing HOPS while providing both Qc for complete SNARE zippering and localized Sec17 apolar loops, the twin driving forces for fusion.

---

Intracellular membrane fusion on the endocytic and exocytic pathways is catalyzed by conserved proteins, including Rab-family GTPases (Hutagalung and Novick, 2011), Rab-binding complexes to tether membranes (Baker and Hughson, 2015), SNARE proteins anchored in each membrane (Jahn and Scheller, 2006), SM (Sec1/Munc18)-family SNARE assembly chaperones (Rizo and Sudhof, 2002), Sec17/SNAP, and the Sec18/NSF AAA-family ATPase (<u>Söllner</u> et al., 1993). Rab GTPases bind their cognate tethering proteins. Tethering allows SNAREs to associate in their parallel, active conformation (Song and Wickner, 2019; Weninger et al., 2003; Zick and Wickner, 2014). SNARE proteins consist of diverse N-domains, 50-60 amino acid SNARE domains with heptad repeat apolar residues, a short juxtamembrane region, and (often) a C-terminal trans-membrane anchor. SNAREs are in 4 conserved families, termed R, Qa, Qb, and Qc according to whether they have a central arginyl or glutaminyl residue instead of their central apolar residue (Fasshauer et al., 1998). Prior to fusion, the R- and Q-SNAREs are on opposite fusion partner membranes (Fukuda et al., 2000). Their SNARE domains alone are inherently random-coil but become alpha-helical as they progressively associate in the N- to C-direction, termed zippering, to form RQaQbQc 4-helical *trans*-SNARE complexes (Sorensen et al., 2006). Zippering is driven by the energy of additional water hydrogen bonding as the SNARE heptad-repeat hydrophobic amino acyl side-chains are buried in the center of the 4-SNARE bundle.

We explore membrane fusion mechanisms through the homotypic fusion of yeast vacuoles (lysosomes). Vacuole fusion has been reconstituted with all purified components (Fukuda et al., 2000; Mima et al., 2008; Stroupe et al., 2009; Zick and Wickner, 2016). The fusion of proteoliposomes of vacuolar mixed lipids (VML) and purified recombinant vacuolar fusion proteins (Mima et al., 2008; Stroupe et al., 2009; Zick and Wickner, 2016) is monitored by a quantitative fluorometric assay (Zucchi and Zick, 2011). Vacuoles and reconstituted proteoliposomes have a GTPase (Ypt7) of the Rab family (Haas et al., 1995), termed simply Rab hereafter. The vacuolar SM protein (Vps33) is a subunit of the hexameric HOPS (**<u>ho</u>**motypic fusion and vacuole **<u>p</u>**rotein **<u>s</u>**orting) protein (Seals et al., 2000; Shvarev et al., 2022), which has Vps 11, 16, 18, 33, 39, and 41 subunits (Wada et al., 1992; Seals et al., 2000). The vacuolar SNAREs (Nichols et al., 1997; Ungermann et al., 1998) are Nyv1 (R), Vam3 (Qa), Vti1 (Qb), and Vam7 (Qc), referred to hereafter as simply R, Qa, Qb, and Qc. Two HOPS subunits, Vps39 and Vps41, have direct affinity for the Rab on each fusion partner membrane, thereby tethering the vacuoles (Brett et al., 2008; Brocker et al., 2012; Hickey and Wickner, 2010). HOPS has direct binding affinity for each of the four SNAREs (Baker et al., 2015; Song et al., 2020), for Sec17 (Song et al., 2021), for Sec18 (Orr and Wickner, 2024), and for phosphoinositides such as PI3P (Stroupe et al., 2006). Parallel grooves on the surface of the SM subunit of HOPS bind the N-terminal end of the R and Qa SNARE domains, positioning them to initiate 4-SNARE assembly (Baker et al., 2015). Early in zippering, the residues of R and Qa which lie in the grooves on the SM HOPS subunit reorient into the center of the nascent 4-helical SNARE complex which may weaken SNARE:HOPS binding. Sec17 can complete the displacement of HOPS from the *trans*-SNARE complex (Collins et al., 2005; Schwartz et al., 2017), and virtually all the *trans*-SNARE complex on vacuoles is associated with Sec17 (Xu et al., 2010). The pre-fusion structure of SNAREs, Sec17, and Sec18 (Song et al., 2021) is modeled on the 20s complex of synaptic SNAREs, NSF, and SNAP (Zhao et al., 2015), but with the SNAREs anchored in *trans*. In this structure, several Sec17/SNAP molecules form a sheath around the 4-SNARE bundle, with the N-terminal apolar loop of each Sec17/SNAP engaging membrane lipids, with ionic bonds between neighboring Sec17 molecules in the sheath (Ma et al., 2016) and between Sec17 and residues on the surface of the 4-SNARE *trans-*complex, and with the Sec17 C-terminal leucyl residues bound to the large Sec18 hexamer. Two models have been envisioned for the assembly of this structure. In one model, the SNAREs first assemble, then several Sec17 molecules bind to the assembled SNAREs and serve as receptor to bind Sec18. In this model, Sec18 would have no effect on the EC50 of Sec17 for fusion. A more recent model (Orr and Wickner, 2022) is that several Sec17 molecules bind reversibly to a Sec18 hexamer as a 3Sec17:Sec18 complex which stably associates with membranes by the product of the weak affinities of the N-domain apolar loop of each Sec17 for lipid. After the interdependent membrane binding of Sec17 and Sec18, their complex binds to SNAREs. This model was suggested by the findings that fusion without complete SNARE zippering is highly cooperative for Sec17 concentration in the absence of Sec18 but not in its presence (Orr and Wickner, 2022), suggesting that Sec18 helps gather multiple Sec17s, and that Sec17 and Sec18 are interdependent for their stable membrane association (Orr and Wickner, 2022; Lopes et al., 2025). In accord with these findings, we now report that Sec17 and liposomes cooperate to enhance ATP hydrolysis by Sec18 in the absence of SNAREs.

Though Sec18 ATPase and Sec17 act after membrane fusion, using energy from ATP binding and hydrolysis to drive the disassembly of *cis*-SNARE complex (Söllner et al., 1993; Haas and Wickner, 1996), Sec17 and Sec18 also act before fusion. Schwartz and Merz (2009) showed that C-terminal truncation of the Qc SNARE domain blocks fusion in a manner that is bypassed by added Sec17 and further stimulated by Sec18 (Zick et al., 2015). Sec17 and Sec18 also bypass fusion blockade from any of several causes: C-terminal truncation of any of the Q-SNAREs (Song et al., 2021), swap of the juxtamembrane domains between R and Qa (Orr et al., 2022), having Qb as the sole membrane anchored Q-SNARE (Wickner et al., 2023), or replacement of the Qa heptad repeat apolar residues with Gly, Ala, or Ser (Song et al., 2021). Even with wild-type SNAREs and unimpeded zippering, Sec17 and Sec18 restore fusion which had been impeded by stiff fatty acyl lipid composition (Song et al., 2024), and Sec18 is essential for fusion with limiting HOPS levels (Orr and Wickner, 2024). Sec17 and Sec18 also stimulate fusion under even the most favorable fusion conditions (Song et al., 2017). Slow fusion can be driven by either SNARE zippering (without Sec17 or Sec18) or by Sec17 and Sec18 (without energy from completion of SNARE zippering), but rapid fusion needs both. The apolarity of the small loop in the Sec17 membrane-proximal N-domain is critical for Sec17 promotion of fusion (Song et al., 2021) and may act as a wedge into the tightly apposed bilayers.

Sec18 can lower the EC50 for the Qc-SNARE for fusion as much as 100-fold (Orr and Wickner, 2024) yet has no effect on the EC50 for the Qb-SNARE. The mechanism of this SNARE-specific Sec18 action has been unclear. We now report that Qc associates with Sec17 and Sec18 on membranes. Though ATP hydrolysis is not needed for membrane fusion (Zick et al., 2015), ATP hydrolysis by Sec18 is a sensitive measure of Sec18 associations with SNAREs, Sec17, and membranes (Barnard et al., 1997; Cipriano et al., 2013; Winter et al., 2009; Barnard et al., 2017).

We find that this Sec18 activation does not require classical RQaQbQc bundles, anchored in *trans* or in *cis*, but needs only Qc, Sec17, and membrane lipid. PI3P localizes Qc to the membrane surface where it can bind the Sec17:Sec18 complex. Sec18:Sec17:Qc:membrane association, assayed by Sec18 activation, is promoted by specific associations: the binding of the Qc PX domain to phosphatidylinositol-3-phosphate (PI3P) at the membrane (Cheever et al., 2001), the recognition of the Qc SNARE domain by Sec17, Sec17 binding to membranes by its N-domain apolar loop (Winter et al., 2009; Zick et al., 2015), and Sec17 binding Sec18 by its C-terminal leucyl residues (Schwartz and Merz, 2009; Barnard et al., 2017). This Qc activation of Sec18 is in accord with the specific association of the Qc SNARE domain in the synaptic SNARE complex with NSF (White et al., 2018). These same interactions between Qc, membranes, Sec17 and Sec18 stimulate membrane fusion. Sec18 may lower the EC50 of Qc for fusion (Orr and Wickner, 2024) by delivering Qc and Sec17 to the rest of the membrane fusion apparatus.

## RESULTS

### Binding relationships between Sec17, SNAREs, and liposomes

To explore SNARE associations, proteoliposomes without proteins or bearing one or several anchored SNAREs were incubated with Sec17, then recovered after floatation up a density gradient and analyzed by immunoblot for bound Sec17. The low levels of Sec17 bound to proteoliposomes with R, Qa, or Qb were comparable to the Sec17 bound to liposomes which bore no protein (Figure 1A, lanes 1-4).

**Figure 1.**
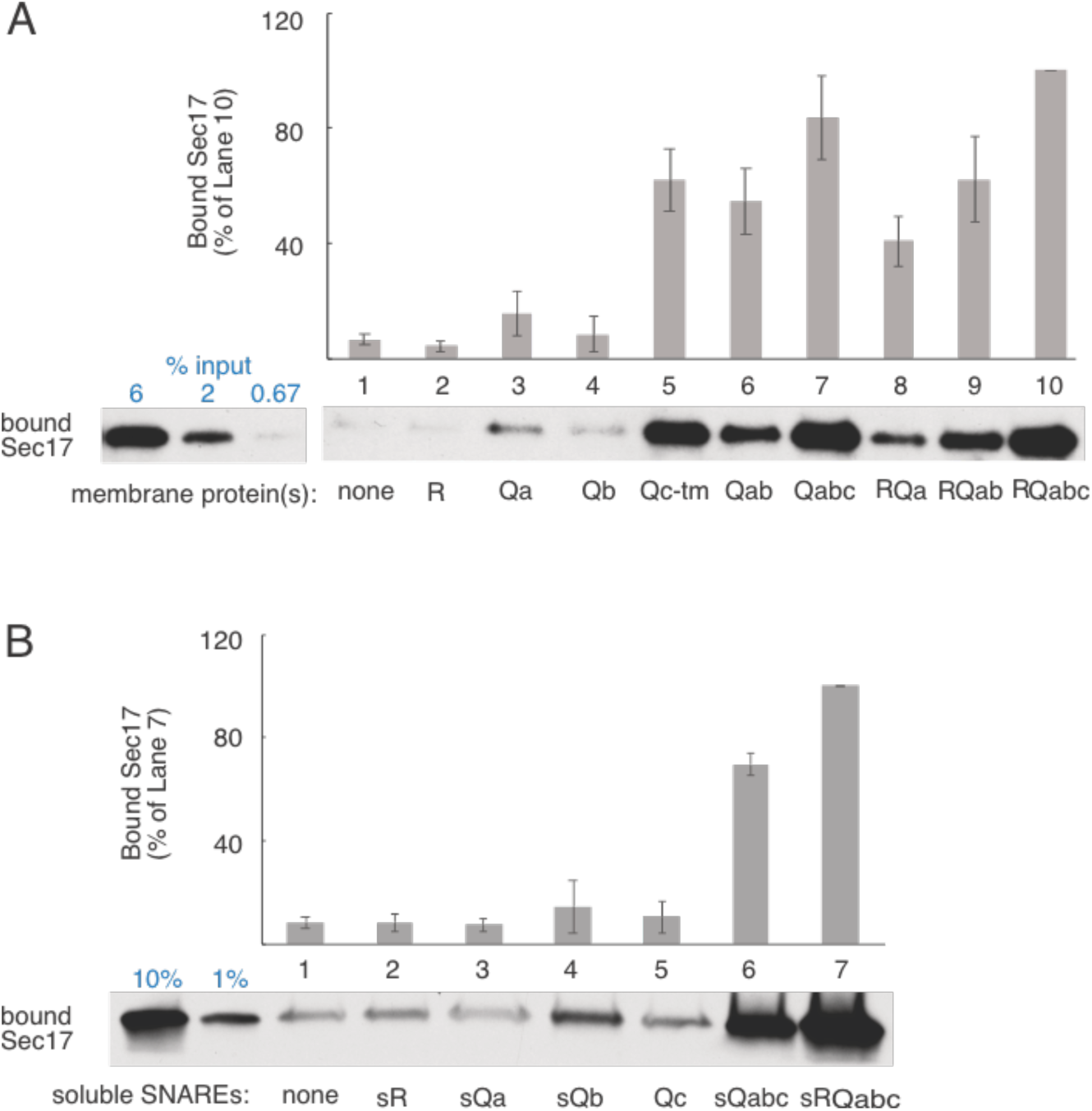
Sec17 binds preferentially to membrane-bound Qc. VML liposomes were prepared for binding assays as described in Materials and Methods with no SNAREs, single membrane bound SNAREs, or combinations of the wild-type SNAREs as indicated. (A) Binding assays were conducted as described in Methods with (A) 1 µM added Sec17 or 1. (B) 2 µM Sec17 and 2 µM soluble SNAREs as indicated.

Substantially more Sec17 bound to proteoliposomes with Qc anchored to membranes by a recombinant C-terminal transmembrane segment, termed Qc-tm (lane 5). Liposomes bearing several SNAREs allowed Sec17 binding (lanes 6-10), even without Qc (lanes 6, 8, and 9), but the addition of Qc gave substantial binding increases (lanes 6 vs 7, and 9 vs 10).

Does Sec17 bind Qc by forming an oligomeric complex of several Sec17 molecules surrounding a coiled coils of several Qc molecules, much as several Sec17 molecules surround RQaQbQc in the 20s structure (Zhao et al., 2015)? When Sec17 is oligomerized, either by association with the 4 soluble-SNAREs (Zick et al., 2015) or with hexameric Sec18 (Orr and Wickner, 2022), it binds membranes with the product of the weak lipid affinities of each of the several Sec17 molecules, converting a very weak lipid affinity of the Sec17 monomer to a very strong liposome affinity of any complex which includes several Sec17 molecules. Neither Qc nor the soluble domains of any of the other three SNAREs, sR, sQa, or sQb, induces Sec17 to oligomerize and thereby bind to liposomes (Figure 1B, lanes 1-5). Sec17 does bind well when it can associate with the three soluble Q-SNAREs (lane 6) or all four soluble SNAREs (lane 7). These findings suggest that Qc alone does not form an oligomer for Sec17 binding.

### Membrane binding of Qc, Sec17, and Sec18

Qc binds to membranes with PI3P (Figure 2, lane 1) and Sec17 binds weakly by its apolar loop to membranes (lane 2). The membrane binding of each is unaltered by the presence of the other (lane 3).

**Figure 2.**
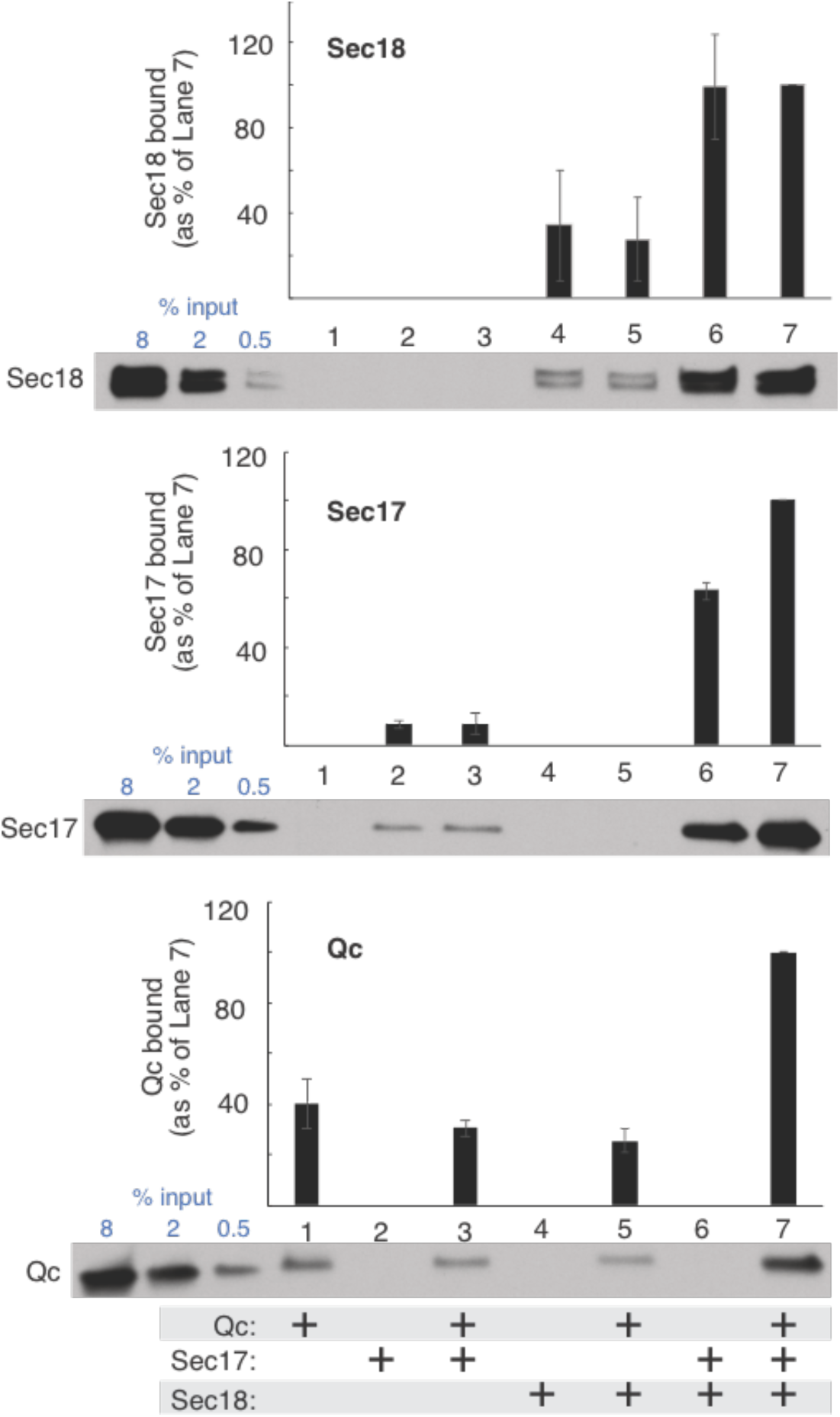
Binding cooperativity between Qc, Sec17 and Sec18. VML liposomes without any membrane proteins were prepared as described for binding assays. Assays were conducted as described with 1 mM Mg:ATPyS added to each reaction, and, where indicated, 1 µM Sec17, 250 nM Sec18, and 2 µM Qc buffer exchanged into 20 mM HEPES-NaOH pH 7.4, 500 mM NaCl, 10% glycerol, 1 mM DTT, 1 mM EDTA.

Sec18 binds to membranes (lane 4) by its affinity for phosphatidic acid (Starr et al., 2016). This Sec18 binding is unaffected by Qc (lane 5). A 3Sec17:Sec18 complex (Orr and Wickner, 2022) strongly enhances both Sec17 and Sec18 membrane binding (lane 6 vs 2 and 4). The 3Sec17:Sec18 complex binds additional Qc to membranes (lane 7 vs 1, 3, 5). Since 4-SNARE coiled-coils assemblies, Sec17, and membranes are known to enhance NSF and SNAP-dependent SNARE complex disassembly (Winter et al., 2009), we asked whether Qc without the other SNAREs could stimulate Sec18 ATPase when it associates with lipid, Sec17, and Sec18.

### Synergy among factors stimulating Sec18 ATPase

We find that Sec18 ATP hydrolysis is exquisitely regulated by interactions between Sec18, Sec17, the Qc-SNARE, and membranes, as illustrated in the scheme of Figure 3A. The number of Qc and Sec17 which associate at the membrane with each other and with Sec18 is not known, but for simplicity only one of each is drawn. There are several direct binding interactions in this scheme: the Qc N-terminal PX domain with its membrane lipid receptor PI3P (Cheever et al., 2001), the N-terminal region of the Qc SNARE domain with Sec18 (White et al., 2018), the SNARE domain of Qc with Sec17 (Marz et al., 2003; Zhao et al., 2015), the N-domain apolar loop of Sec17 with the lipid bilayer (Winter et al., 2009; Zick et al., 2015), and the C-terminal leucyl residues of Sec17 with Sec18 (Barnard et al., 1997; Schwartz and Merz, 2009; Barnard et al., 2017). We asked whether each regulates ATP hydrolysis by Sec18 (Figure 3B and C; T tests of significance are shown in parenthesis in the text for Figure 3B for clarity).

**Figure 3.**
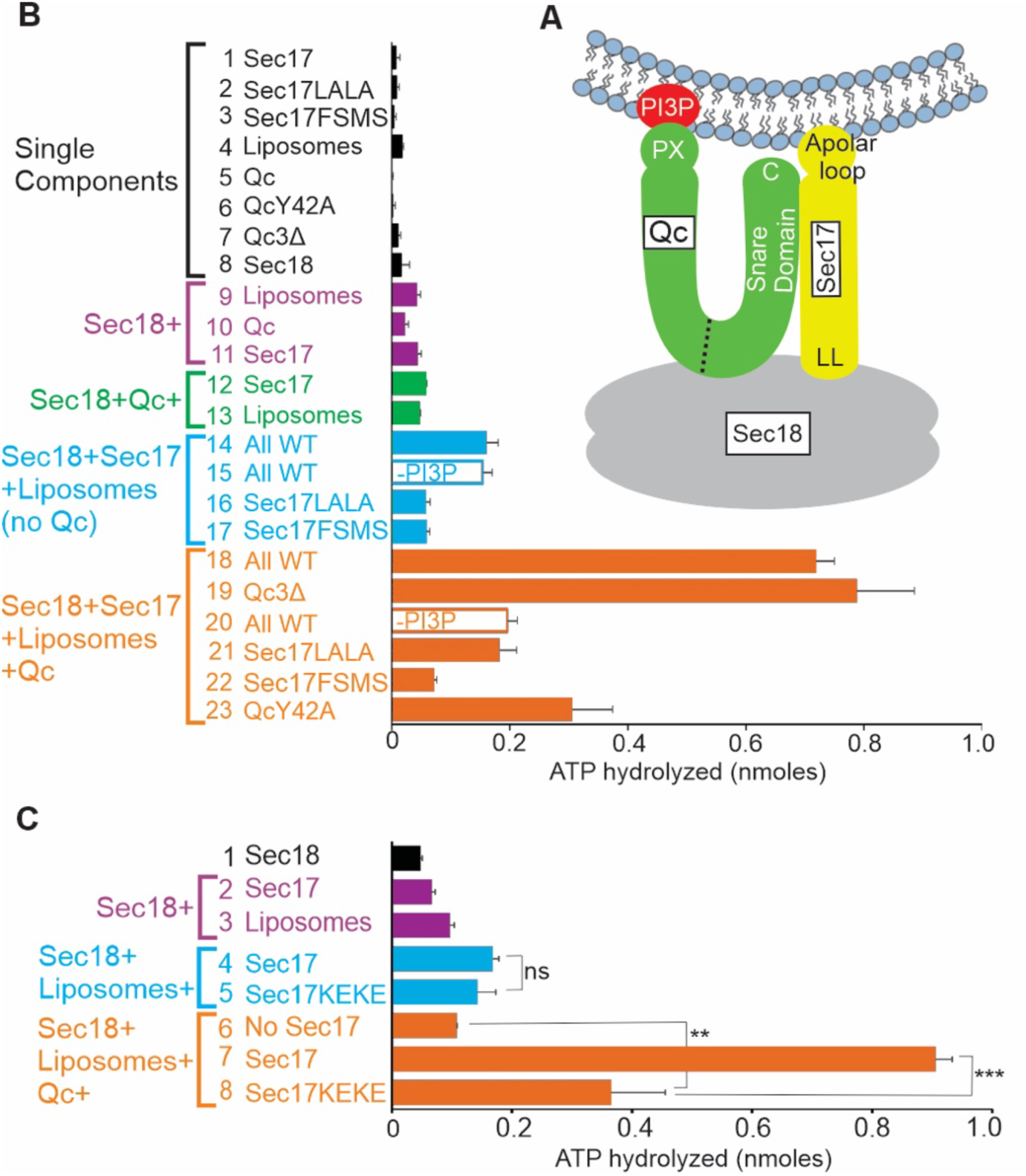
Assaying Sec18 associations by ATP hydrolysis. (A) Working model. Stimulation of Sec18 ATP hydrolysis by membrane-bound Sec17 and Qc requires specific interactions of Qc, Sec17, and Sec18 with each other and with membranes. (B) ATPase assays (20 μl) were performed for 30 min as described in Methods with 0.4 mM VML (vacuolar mimic lipid) liposomes, 50 nM Sec18, 1.5 μM Sec17 or its mutants, 1 μM Qc or its mutants, individually or in combinations as indicated. (C) Sec17 needs its SNARE-binding lysyl residues to stimulate Sec18 ATP hydrolysis when Qc is present. Assays of ATP hydrolysis, as described in Methods, were for 20 minutes and had (where indicated) 0.4 mM liposomes, 100 nM Sec18, the indicated concentrations of Sec17 or Sec17KEKE, and 1 μM Qc.

Sec18 hydrolysis of ATP (Söllner et al., 1993) showed only slight stimulation by either Sec17 (Figure 3B, lanes 11 vs 8_*_) as reported (Morgan et al., 1994) or by protein-free liposomes of vacuolar lipid composition (lanes 9 vs 8_*_) and was not stimulated by Qc SNARE alone (lanes 10 vs 8, ns). Combinations of Sec18, Sec17, liposomes, and Qc caused synergistic stimulations. There was greater stimulation of Sec18 ATP hydrolysis by Sec17 and liposomes (lane 14) than by either Sec17 (lane 14 vs 11_***_) or liposomes (lane 14 vs 9_***_) alone. This relied on the Sec17 C-terminal leucyl resides which bind Sec18 (Barnard et al., 1997; Schwartz and Merz, 2009; Bernard et al., 2017) and are substituted in Sec17L291AL292A, termed Sec17LALA (lane 16, vs wild-type Sec17 in lane 14_***_). Stimulation also relied on the Sec17 N-domain loop apolarity for binding membranes (Winter et al., 2009; Zick et al., 2015), substituted in Sec17F21SM22S, termed Sec17FSMS (lane 17, vs wild-type in lane 14_***_). We have previously reported two independent assays of the interactions of Sec17 with both membranes and Sec18. In functional assays of fusion with blocked SNARE zippering, Sec18 is required for fusion at limiting Sec17 (Orr and Wickner, 2022). In a physical assay of association, Sec17 and Sec18 exhibited interdependent binding to protein-free liposomes in floatation assays (ibid, and Figure 2). The finding that lipids and Sec17 strongly stimulate the ATPase activity of Sec18, and that this stimulation needs the Sec17 binding sites for lipid and Sec18, provides a third and independent indication of Sec18:Sec17:lipid interactions.

### Stimulation by the Qc SNARE

The further presence of Qc-SNARE causes substantially greater stimulation of Sec18 ATP hydrolysis (Figure 3B, lane 18). This ATPase activity relies on Sec17 (lanes 18 vs 13_***_), Qc (lanes 18 vs 14_***_), and liposomes (lanes 18 vs 12_***_). PI3P, a membrane receptor for Qc (Cheever et al., 2001), was needed for Qc stimulation of Sec18 ATPase in the presence of Sec17 and Qc (lanes 18 vs 20_***_) though PI3P omission had no effect on ATP hydrolysis without Qc (lanes 14 vs 15, ns). The Y42A mutation in the Qc PX domain, which ablates the Qc affinity for PI3P (Cheever et al., 2001), reduced its stimulation of Sec18 ATPase (lanes 18 vs 23_***_). In contrast, Qc stimulation of Sec18 ATPase in the presence of lipids and Sec17 does not require the C-terminal portion of the SNARE domain (lane 18 vs 19 ns). Proteins which contribute to optimal Sec18 ATPase have direct membrane affinity: Sec17 by its N-domain apolar loop which is ablated by the FSMS mutation and Qc by the affinity of its PX domain for PI3P which is lost without this lipid or when Qc binding was inactivated by the Y42A mutation. Sec18 binds membranes by its affinity for phosphatidic acid (Starr et al., 2016) as well as through its bound Sec17 (Orr and Wickner, 2022).

Sec17 binds Sec18 by the Sec17 C-terminal leucyl residues and binds membranes by the Sec17 apolar loop (Winter et al., 2009; Zick et al., 2015; Orr and Wickner, 2022). Qc may engage the Sec17:Sec18 complex by direct affinity of its SNARE domain for Sec17 (Zhou et al., 2015; also see Figures 1, 3C and [below] 5) and for Sec18 (White et al., 2018), while the Qc N-terminal PX domain engages membrane PI3P (Cheever et al., 2001). The assembly of membrane, Qc, Sec17, and Sec18 has at least two functional redundancies for stimulating Sec18. There is more ATP hydrolysis when Qc binding to membrane PI3P is lost through the Y42A mutation than when Qc is altogether absent (Figure 3B, lane 23 vs 14_*_), suggesting that Qc can functionally engage through only binding Sec17 and Sec18. Similarly, when the Sec17 binding to Sec18 is weakened by the Sec17LALA mutation, Sec18 ATP hydrolysis is reduced but remains greater than seen without Sec17 (lane 21 vs 13_***_) or without Qc (lane 21 vs 16_***_).

### Sec17:Qc SNARE domain recognition

Sec17 is adjacent to the SNARE domains of Qc and the other SNAREs in the classic 20s structure (Zhao et al., 2015). The ionic bonds between Sec17 and SNARE domains are weakened by substitution of acidic amino acyl residues for basic lysyl residues in the Sec17 mutant Sec17K159EK163E (Marz et al., 2003, Schwartz et al., 2017), termed Sec17KEKE. Sec17KEKE retains its overall structure since, in the absence of Qc, Sec17KEKE stimulation of Sec18 ATP hydrolysis is comparable to that from wild-type Sec17 (Figure 3C, lane 4 vs 5). In the presence of Qc, Sec17KEKE stimulates less than wild-type Sec17 (lanes 7 vs 8) though more than seen in the absence of Sec17 (lanes 8 vs 6). Though the Rab Ypt7 is required for fusion of vacuoles (Haas et al., 1995; Seeley et al., 2002) or of vacuole-derived proteoliposomes (Stroupe et al., 2009; Mima and Wickner, 2009), it does not affect the hydrolysis of ATP by Sec18 (Supplementary Figure S1).

### Membrane-anchored SNAREs

Since Qc must bind membranes to stimulate the Sec18 ATPase, we compared the efficacy of soluble, wild-type Qc with Qc bearing a recombinant C-terminal transmembrane anchor (Xu and Wickner, 2012), termed Qc-tm, in assays of Sec18 ATPase activity with Sec17. ATP hydrolysis with Qc from 0 to 1000 nM (Figure 4A, lanes 6-10) with protein-free liposomes were compared to a reaction with liposomes bearing Qc-tm (lane 13). Incubations with equal stimulations of ATP hydrolysis by Sec18 (lanes 8 and 13) were analyzed by immunoblot and had the same concentrations of Qc (Figure 4A, top). Wild-type Qc, bound to membranes by its affinity for PI3P, has a similar stimulatory activity as transmembrane-anchored Qc.

**Figure 4.**
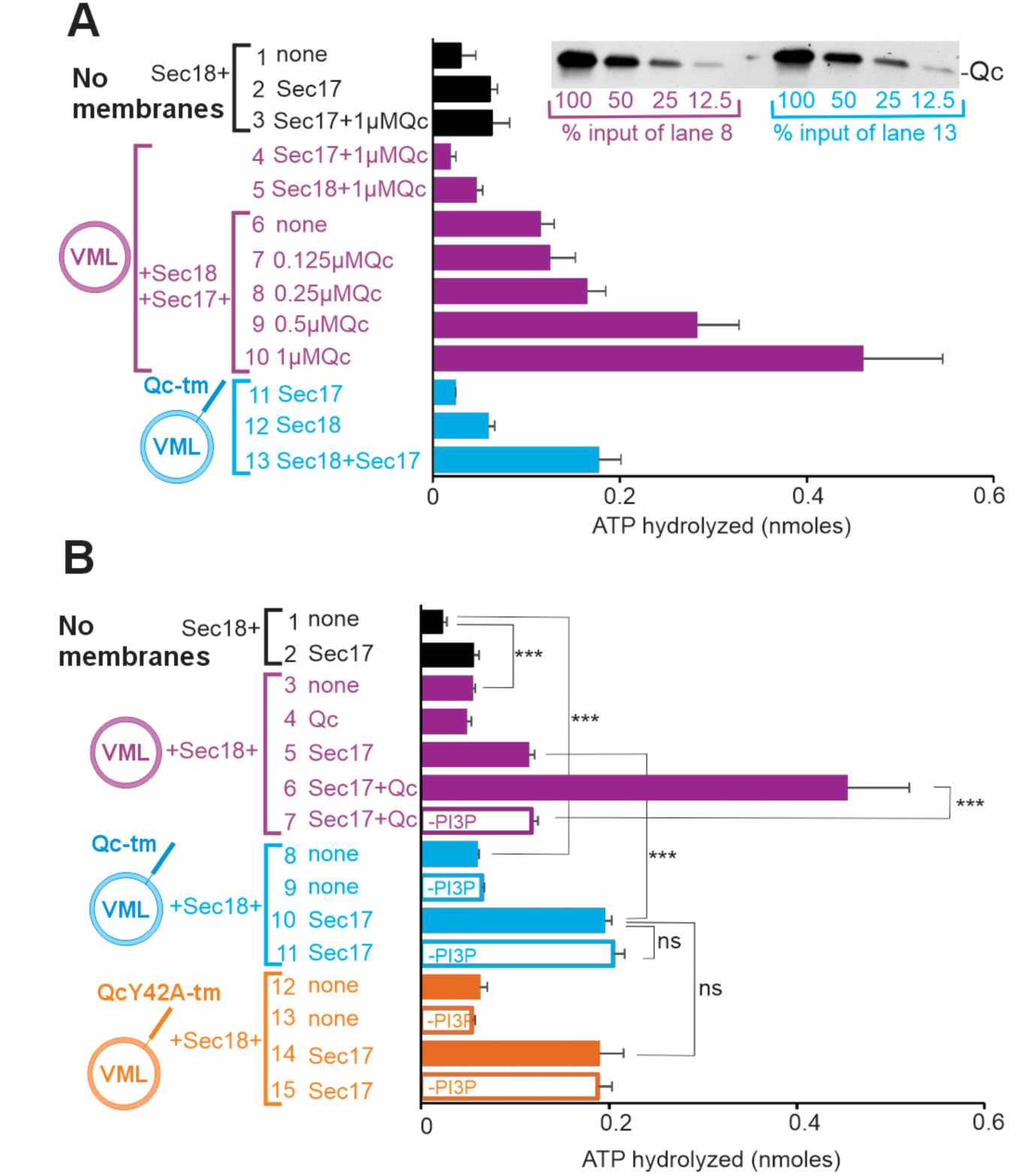
Stimulation of ATP hydrolysis by Qc-tm is as effective as Qc. (A) With the same amount of Qc in the incubation, there is equal Sec18 stimulation by Qc or Qc-tm. ATPase assays (10 minutes) were performed with 0.4 mM VML lipid, 1.5 μM Sec17, 50 nM Sec18, and 1, 0.5, 0.25, 0.125 or 0 μM Qc as indicated. Top, immunoblot; left four lanes are 2-fold dilutions of tube 8, the right four lanes are 2-fold dilutions of tube 13. (B) Anchored Qc-tm or QcY42A-tm doesn’t need PI3P. ATPase assays (10 minutes, as described in Methods) had (where indicated) 0.4 mM membrane lipids (either complete vacuolar mixed lipids [VML] or with PI3P omitted), 1.5 μM Sec17, 50 nM Sec18, and 2 μM Qc.

To determine whether the only role of PI3P in support of Sec18 ATPase is to bind Qc to the membrane, we assayed Sec18 ATP hydrolysis in the presence or absence of Sec17 and either no liposomes (Figure 4B, black), liposomes without proteins (purple), liposomes bearing Qc-tm (blue) or QcY42A-tm to ablate affinity for PI3P (orange). Liposomes without membrane proteins, with Qc-tm, or with QcY42A-tm were also prepared without PI3P. Without Sec17, liposomes with Qc-tm only stimulated Sec18 to the same limited extent as liposomes without Qc-tm (Figure 4B, lanes 3 and 8 vs 1), but Qc-tm stimulated fusion in the presence of Sec17 (lanes 10 vs 5). PI3P is needed as the membrane receptor for wild-type Qc to stimulate Sec18 ATPase (lanes 6 vs 7) but is not needed with anchored Qc-tm (lanes 10 vs 11). Though the ablation of PI3P affinity through the Y42A mutation in Qc reduces its capacity to bind the membranes and stimulate Sec18 (Figure 3B, lanes 18 vs 23), the Y42A mutation has no effect when the Qc is tm-anchored (Figure 4B, lanes 10 vs 14). The sole function of PI3P for stimulating Sec18 ATPase is therefore as a membrane binding site for Qc.

### Qc-SNARE regions which activate Sec18

Which parts of the Qc-SNARE activate Sec18? Proteoliposomes were prepared without protein, with Qc-tm, with Qc-tm deleted for its N-terminal PX domain and part of the remaining N-domain (Xu and Wickner, 2012) (termed here QcΔPX-tm), or with Qc-tm deleted for its entire N-domain, leaving only the anchored SNARE domain (QcSD-tm). Each [proteo]liposome was incubated with Sec18. Without Sec17, there was little enhancement of the Sec18 ATPase by liposomes bearing any of the 3 forms of Qc beyond that seen with protein-free liposomes (Figure 5A, lane 5, 8, 11, 14). In the presence of Sec17, ATP hydrolysis was enhanced equally by Qc-tm or QcΔPX-tm (lanes 6 vs 9 or 12), while QcSD-tm (lane 15) gave maximal stimulation. Thus, the anchored SNARE domain of Qc interacts with Sec17 and Sec18 for maximal ATPase activity, modulated by the Qc N-domain.

**Figure 5.**
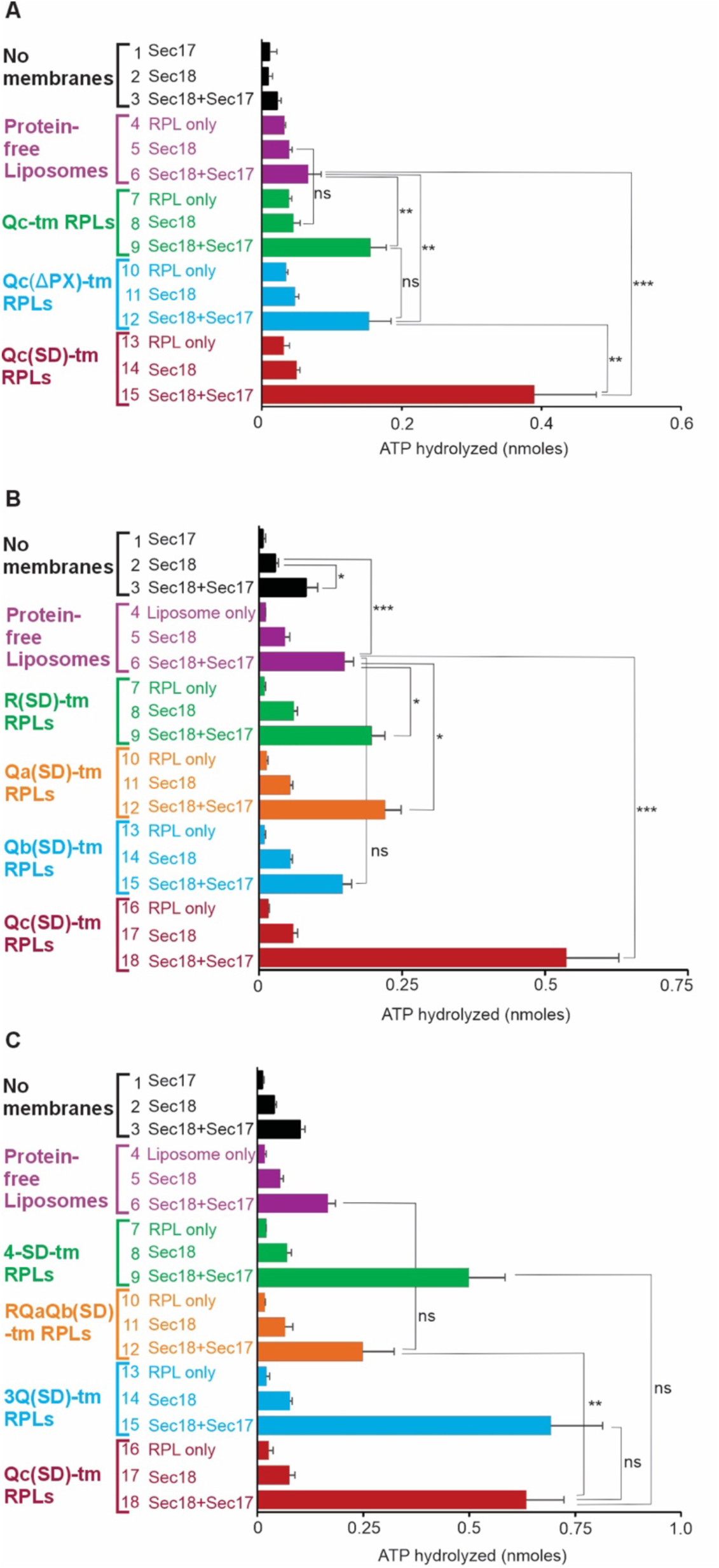
Potency of the Qc SNARE Domain. (A) QcΔPX-tm or QcSD-tm are as effective as Qc-tm in supporting Sec1718 ATPase. ATPase assays (10 minutes, as in Methods) had (where indicated) proteoliposomes with 0.4 mM lipids, 50 nM Sec18, and 500 nM Sec17. Proteoliposomes had no protein (purple) or a 1:2000 molar ratio to lipid of Qc-tm (green), QcΔPX-tm (blue), or QcSNAREDomain-tm (red) as indicated. (B) QcSD-tm is far more effective than R-, Qa-, or Qb-SD-tm. ATP hydrolysis assays (40 minutes, as in Methods) contained 3 μM Sec17, 50 nM Sec18, and no proteoliposomes (lane 3), protein-free proteoliposomes (lane 6) or proteoliposomes bearing either R, Qa, Qb, or Qc SNARE Domain with a C-terminal transmembrane anchor (lanes 9, 12, 15, 18). As controls, lanes 5, 8, 11, 14, and 17 had the indicated proteoliposomes with 50 nM Sec18 but no Sec17, while lanes 4, 7, 10, 13, and 16 had only proteoliposomes without either Sec17 or Sec18. (C) QcSD-tm is more active than RQaQbSD-tm. ATPase assays (Methods, 40minutes) had (where indicated) 50 nM Sec18, 3 μM Sec17, and proteoliposomes without membrane proteins (lanes 4-6), all 4 tm-anchored SNARE domains (7-9), and R, Qa, and Qb tm-anchored SNARE domains (10-12), all three tm-anchored Q SNARE domains (13-15), or QcSD-tm (16-18).

### Comparing anchored SNARE domains

Is the Qc SNARE domain uniquely active as a Sec17 ligand for activating Sec18 ATPase, or can other SNARE domains, singly or in combination, fulfill this function as efficiently? Proteoliposomes of vacuolar lipid composition were prepared without protein, with the R, Qa, Qb, or Qc-tm individual SNARE domains without their N-domains but with a C-terminal membrane anchor, or with several of these membrane-anchored SNARE domains. The low ATPase activity of Sec18 alone (Figure 5B, lane 2) is modestly enhanced by simply Sec17 (lane 3), less so by liposomes without Sec17 irrespective of their anchored SNARE(s) (lanes 2 vs 5, 8, 11, 14, or 17). Sec18 was further stimulated by the combination of Sec17 and liposomes (lane 6), with modest additional activity from anchored R- or Qa-SNARE domain (lanes 9 and 12). There was no additional stimulation by anchored Qb SNARE domain (lane 15). The greatest stimulation of the Sec18 ATPase was by anchored Qc SNARE domain (lane 18). Sec18 ATP hydrolysis stimulated by membrane-anchored Qc SNARE domain with Sec17 was not enhanced by the other anchored Q-SNARE domains (Figure 5C, lanes 15 vs 18), even with the anchored R-SNARE domain as well (Figure 5C, lanes 9 vs 18). Strikingly, the combination of anchored R, Qa, and Qb SNARE domains without Qc-tm gave less stimulation than Qc-tm alone (lanes 12 vs 18) and was barely above the signal with protein-free liposomes (lanes 12 vs 6) even though R, Qa, and Qb form a stable 3-SNARE complex (Song et al, 2020). Multiple anchored SNARE domains are not sufficient for full stimulation of Sec18 ATPase unless they contain the Qc-SNARE.

### The same proteins and their interactions govern HOPS-dependent fusion and Sec18 ATP hydrolysis

To assay how Sec18, Sec17, Qc, and membranes with PI3P regulate fusion, proteoliposomes bearing Rab, R, and lumenal biotinylated phycoerythrin were mixed with those bearing Rab, Qa and lumenal Cy5-streptavidin. Mixed soluble components were added: HOPS, soluble Qb (without a transmembrane anchor), Qc, Sec17 where indicated, Sec18 where indicated, and MgATP. Fusion was assayed as the mixing of the proteoliposome lumenal compartments, allowing the two fluorescent proteins to bind and generate a strong FRET signal (Zucchi and Zick, 2011). With neither Sec17 nor Sec18, there is slow fusion which is driven by complete SNARE zippering (Figure 6A, lane 1). There is minimal stimulation by 100 nM Sec17 (Figure 6A, lane 2). Mutant forms of Sec17 were used to selectively disable the Sec17 interactions with either the membrane, SNAREs, or Sec18. Fusion was statistically equivalent with either wild-type Sec17 (lane 2) or when the apolarity of the Sec17 N-domain loop was ablated by the FSMS mutation (lane 3), with Sec17LALA which lacks the Sec17 binding site for Sec18 (lane 4), or when the Sec17 binding site for SNAREs was weakened by the KEKE mutation (lane 5). Without Sec17, there was little stimulation by Sec18 (lane 6 vs 1). The combination of Sec17 and Sec18 (Figure 6A, lane 7) gave a substantial stimulation of fusion (compare lane 7 with lanes 1, 2, and 6). This maximal fusion (lane 7) is inhibited by Sec17 mutants which interfere with its interaction with membrane lipids (lane 8), with Sec18 (lane 9), or with SNAREs (lane 10). The inhibition from loss of Sec17 affinity for lipids or SNAREs is almost to the level of fusion seen without Sec17 at all (compare lanes 6 with 8 and 10). With Sec17LALA, which lacks affinity for Sec18, Sec18 has no effect on fusion (compare lanes 4 and 9). Supplemental Figure S2, which contains examples of the kinetics of fusion (panels A and B), also shows that QcY42A, without affinity for membrane PI3P (Cheever et al., 2001), does not support fusion at all, even with Sec17 or Sec18 (panel C). Thus, each protein (Sec17, Sec18, and Qc) and each interactive domain needed for optimal ATP hydrolysis, as seen in Figure 3, also underlie optimal fusion.

**Figure 6.**
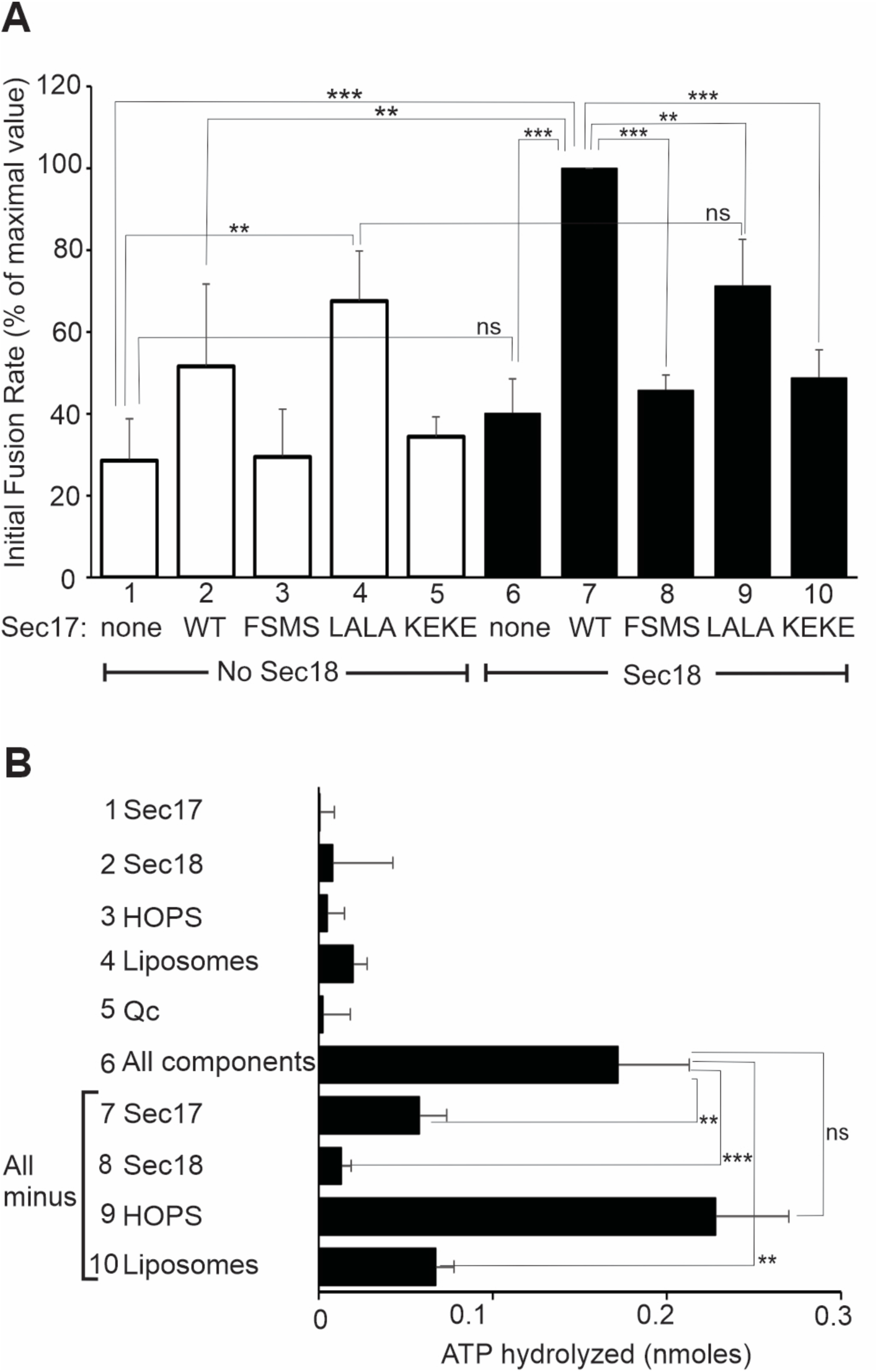
Sec17, Sec18, membranes, and their interactions affect membrane fusion as well as ATP hydrolysis. (A) Sec17 and Sec18 stimulate membrane fusion. Fusion was assayed (Orr and Wickner, 2024) as the FRET between biotin-phycoerythrin and Cy5-streptavidin, each initially entrapped in separate proteoliposome populations (Zucchi and Zick, 2011). Assays (20 μl) had proteoliposomes of vacuolar mixed lipid composition with Rab and R or with Rab and Qa (1:8,000 molar ratio of Rab to lipids and 1:16,000 molar ratio of SNARE to lipid). Each assay was in the presence of excess nonfluorescent streptavidin to block any FRET signal from proteoliposome lysis. Mixed proteoliposomes were initially incubated with GTP and EDTA to strip bound guanine nucleotide, then given excess MgCl_2_. Proteoliposomes and mixed soluble proteins (50 nM HOPS, 100 nM Sec17 where indicated, 50 nM Sec18 where indicated, 1 μM sQb, 20 nM Qc, and 1 mM MgATP) were preincubated in separate wells for 10min at 27°C, then mixed with a multichannel pipettor. Assays were performed in 384 well plates (Item 4514, Corning, Kennebunk, ME) in a Gemini XPS plate reader (Molecular Devices, Sunnyvale, CA). Average and standard deviations of the initial rates are shown for triplicate experiments. Fusion was performed either without Sec18 (open bars) or with 50 nM Sec18 (filled bars) and either without Sec17 or with 100 nM Sec17, either wild-type, FSMS, LALA, or KEKE. (B) HOPS, the essential catalyst of tethering and *trans*-SNARE assembly for fusion, is not needed for ATP hydrolysis. Proteoliposomes were prepared for fusion assays as described in Methods, but each internal fluorescent protein stock had first been dialyzed twice against a 200-fold volume excess of Rb150 to remove phosphate. Fusion incubations, as described above, had Rab/R and Rab/QaQb proteoliposomes (333 μM lipids, 1:16,000 molar ratios of SNAREs:lipids, 1:8,000 of Rab:lipids) and 100 nM HOPS, 300 nM Sec17, 50 nM Sec18, 1mM MgATP, and 1 μM Qc except where omitted as noted. Incubations (20 min, 23°C, 0.5 ml conical plastic tube) were followed by assay of inorganic phosphate as described in Methods.

## ATP hydrolysis by Sec18 is not affected by HOPS

HOPS is essential for fusion, but does it regulate ATP hydrolysis? Fusion components were mixed and incubated under fusion assay conditions. EDTA was added to terminate ATP hydrolysis, and each sample was assayed for the inorganic phosphate released from ATP. Each separate reaction component only showed minimal phosphate contamination or generation (Figure 6D, lanes 1-5), but together there was substantial ATP hydrolysis (lane 6). In assays where single reaction components were omitted, there was no effect of omitting HOPS (lane 9), but ATP hydrolysis was diminished by omission of Sec18, Sec17, or proteoliposomes (lane 6 vs 7, 8, 10). Since HOPS is required for *trans*-SNARE complex assembly, this confirms that HOPS, and presumably its receptor Rab and the *trans*-complex it assembles between R and Qa SNAREs, are not needed for the activation of Sec18 ATPase. The initial HOPS interactions with Rab and the R and Qa SNAREs are distinct from the initial interactions of Sec18, Sec17, Qc, and membranes.

## DISCUSSION

Our studies are summarized in a working model (Figure 7). Membrane-bound Qc (Figure 7, part i) binds to the membrane-bound complex of Sec17 and Sec18 (ii) to form a complex of membrane-bound Sec18, Sec17, and Qc (iii). Independently, HOPS binds to the Rab on each membrane for tethering, then catalyzes the initial assembly of R and Qa on apposed membranes to form the HOPS-RQa *trans* complex, which binds Qb (iv). Without Qc, RQaQb zippering does not proceed to fusion. The Sec18:Sec17:Qc complex binds the HOPS:R:QaQb complex by some combination of the affinities of Qc for the other SNAREs and for HOPS and of Sec17 for HOPS and for the entire SNARE complex. HOPS is displaced during this assembly, yielding the Sec18:Sec17*:trans*-4-SNARE prefusion complex (v). While the stoichiometry and structures of each of these complexes is not known, each is supported by prior and current studies.

**Figure 7.**
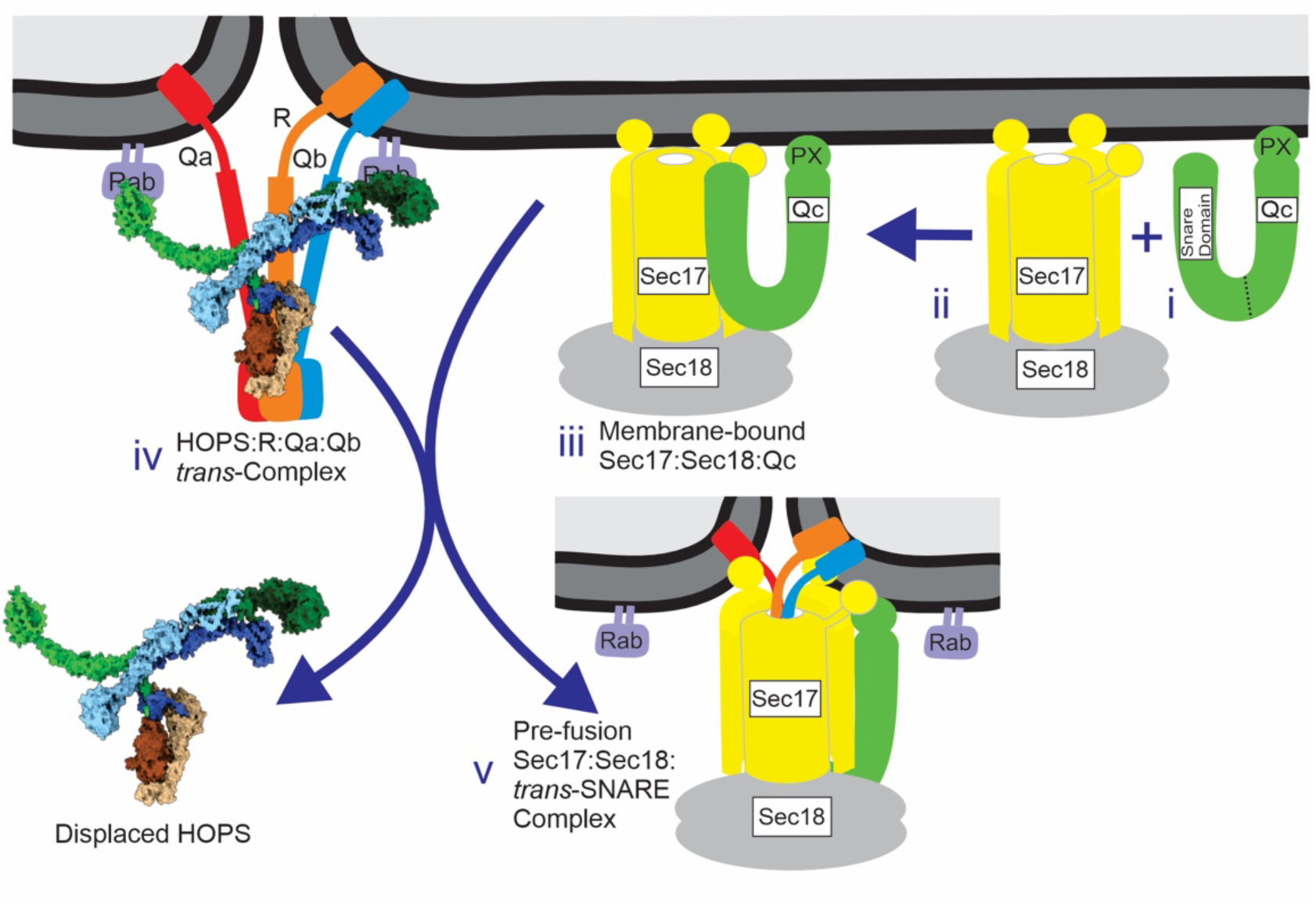
Current working model of the Sec17:Sec18:Qc complex and the HOPS:R:Qa:Qb complex combining while displacing HOPS, forming the prefusion complex. There is evidence in support of each intermediate: i) Qc is membrane-bound by the affinity of its N-terminal PX domain for PI3P (Cheever et al., 2001), ii) Sec18 and Sec17 assemble into a 3Sec17:Sec18 complex which binds to lipid by the Sec17 apolar loops (Orr and Wickner, 2022) and thereby activating the Sec18 ATPase (Figure 3), iii) Qc binds to Sec17:Sec18 at the membrane (Figure 2) by its affinity for Sec17 (Figure 1A) and for Sec18/NSF (White et al., 2018), further activating ATP hydrolysis (Figure 3B), iv) HOPS catalyzes the assembly of R:QaQb rapid-fusion intermediates (Song et al., 2020), v) HOPS and Sec17 never associate with the same SNARE complexes (Collins et al., 2005), and Sec17 can displace HOPS from SNAREs (Collins et al., 2005; Schwartz et al., 2017).

**Membrane-bound Qc** (i) The Qc-SNARE binds membranes by the affinity of its N-terminal PX domain for PI3P (Cheever et al., 2001). Qc is comprised of an N-terminal PX domain, a helical coils region, a hinge region, and a C-terminal SNARE domain. The affinity of the Qc PX domain for PI3P (Cheever et al., 2001) is needed for wild-type Qc binding to membranes and optimal activation of Sec18 (Figure 3B), but PI3P is not needed for Sec18 stimulation with Qc-tm which is integrally anchored (Figure 4). The PX domain is folded back on the SNARE domain to bind the membrane adjacent to the C-terminus of Qc and other Q-SNAREs. To support fusion, each SNARE domain must remain oriented with its C-terminus near the membrane to engage Sec17 and the other 3 SNAREs as shown in the 20s structure (Zhou et al., 2015).

**Membrane-bound complex of Sec17 and Sec18** (ii) Independent of Qc, Sec18 and several Sec17 assemble and undergo interdependent binding to membranes (Orr and Wickner, 2022) by the product of the affinity of the apolar loop of each of the several Sec18-bound Sec17 molecules for lipid (ibid). Membrane-bound Sec18:3Sec17 stimulates fusion at low Sec17 levels and eliminates the cooperativity of fusion on Sec17 concentration (ibid). Lipid and Sec17/SNAP can cooperate to enhance 4-SNARE complex disassembly by Sec18/NSF (Winter et al., 2009) and they also stimulate Sec18 ATP hydrolysis without SNAREs (Figure 4A, lane 1 vs 6).

**Sec17:Sec18:Qc complex on membranes** (iii) Qc has direct physical and functional interactions with Qa and Qb in the 3Q or 4-SNARE bundle (Sutton et al., 1998), with HOPS (Song et al, 2020), PI3P (Cheever et al., 2001), Sec17 (Figure 1A), and Sec18 (White et al., 2018). Qc association with the 3Sec17:Sec18 complex (Figure 2A) to form Qc:3Sec17:Sec18 may exploit the high affinity of Sec17 for Qc (Figure 1A). More Qc binds to membranes with Sec17 and Sec18 (Figure 2, lane 7) than with either Sec17 or Sec18 alone (lanes 3 and 5). The anchored Qc SNARE domain in Qc:3Sec17:Sec18 activates the Sec18 ATPase (Figure 5A, lane 15). The C-terminal third of this SNARE domain is not needed (Figure 3B, lane 19), in accord with only the N-terminal portion of the synaptic SNARE complex being engaged with NSF (White et al., 2018). Optimal Qc:Sec17 interaction on membranes may rely on Qc being in a loop-like conformation with the C-terminus of its SNARE domain adjacent to its N-terminal PX binding site for PI3P. In this conformation, the SNARE domain of membrane-bound Qc can bind Sec17 and be properly presented to Sec18. The affinity of the Qc PX domain for PI3P (Cheever et al., 2001) is needed for wild-type Qc binding to membranes and activation of Sec18, and Qc needs its PI3P recognition to support fusion (Figure 6A), but PI3P is not needed for Sec18 ATPase stimulation with Qc-tm which is integrally anchored. Three recognition surfaces on Sec17 are critical, both for Sec17 stimulation of fusion in the presence of Sec18 and for its activation of the Sec18 ATPase. Both activities are diminished by either the Sec17FSMS abrogation of N-loop apolarity (Figure 3B and 6A), by diminished SNARE affinity of Sec17KEKE (Figure 3C and 6A), and by diminished Sec18 affinity of Sec17LALA (Figure 3B and 6A).

**HOPS:RQaQb complex** (iv) Prior to fusion, HOPS tethers two membranes by binding to the Rab on each. The SM subunit of HOPS (Vps33) binds the R and Qa SNAREs to initiate their zippering (Baker et al., 2015; Orr et al., 2017; Jiao et al., 2018), further engaging the Qb SNARE to form the initial HOPS:RQaQb *trans*-SNARE complex. The HOPS:RQaQb trans-complex between tethered membranes has been reconstituted with proteoliposomes and supports strikingly rapid fusion upon engaging Qc (Song et al., 2020).

**Assembly of the prefusion complex** (v) 3Sec17:Sec18:Qc association with the HOPS:trans-RQaQb complex may exploit HOPS recognitions of Qc (Song et al., 2020; Zick and Wickner, 2013), Sec17 (Orr and Wickner, 2022), and Sec18 (Orr and Wickner, 2024), Qc engagement with the other SNAREs, and Sec17 affinity for a complex of RQaQb (Figure 1, lane 9). As Qc enters and completes the *trans*-SNARE complex, Sec18 and Sec17 may displace HOPS (Collins et al., 2005; Schwartz et al., 2017).

The SNAREs on isolated vacuoles are in complex with either HOPS or with Sec17 but not with both (Collins et al., 2005), suggesting that HOPS displacement by Sec17:Sec18:Qc is efficient. Since Sec18 dramatically lowers the EC50 of Qc for fusion (Orr and Wickner, 2024), Qc either associates with Sec17:Sec18 as a prelude to joining the other SNAREs (Figure 7) or Sec17:Sec18 joins the HOPS:RQaQb complex and enhances its affinity for Qc. Further studies are needed to test these alternative models.

## Materials and Methods

### Reagents

Lipids were from Avanti Polar Lipids (Alabaster, AL), Echelon Biosciences (Salt Lake City, UT), Sigma-Aldrich (Burlington, MA) and Invitrogen (Eugene, OR). Biotinylated R-phycoerythrin and glycerol were from Invitrogen, and Cy5-streptavidin from SeraCare (Milton, MA).

Dialysis tubing was from Repligen (Waltham, MA) and BioBeads from Bio-Rad (Hercules, CA). n-Octyl-β-D-glucopyranoside (ß-octylglucoside) was from Anatrace (Maumee, OH). HEPES, Thesit, Histodenz, ATP, ATPyS, IPTG, and defatted BSA (bovine serum albumin) were from Sigma-Aldrich. Dithiothreitol (DTT) was from Research Products (Mt. Prospect, IL).

### Proteins

GST-Vam3, Vti1, and Nyv1 were purified as described (Mima et al., 2008). Vti1 and Nyv1 were buffer exchanged into Rb150 (20 mM HEPES-NaOH, pH 7.4, 150 mM NaCl, 10% glycerol) + 1% ß-octylglucoside (Zucchi and Zick, 2011).

The Qc-SNARE Vam7 (Schwartz and Merz, 2009) was purified as described (Starai et al., 2007) with modifications. The plasmid encoding Qc (Schwartz and Merz, 2009) was used as template DNA to make the QcY42A point mutant using the primers from Fratti et al. (2007). For the wild-type Qc and the Y42A mutant, plasmid DNA was transformed into C43(DE3)-RP cells and plated on LB + kanamycin and chloramphenicol. A clone was grown in 200 ml LB (L Broth) + Kan/Cam overnight with shaking at 37°C. In the morning, 60 ml was transferred into each of three 4 liter baffled flasks containing 2 liters of Terrific Broth, TB salts, Kan, and Cam, and shaken at 37°C to an OD of 2.0. Expression was induced with 1 mM IPTG and cultures grown for 4 hours with shaking.

Cells were sedimented (JLA10.5 rotor, 5,000 rpm, 5 min, 23°C) then resuspended in 80 ml of Buffer A (20 mM HEPES-NaOH, pH 8.0, 0.5 M NaCl, 0.1% Triton-X100, 1 mM EDTA) and French pressed twice. Lysates were centrifuged in a Beckman 60Ti fixed angle rotor (50,000 rpm, 30 min, 2°C). Supernatents were divided between two Falcon tubes, each with 14 ml chitin resin (NEB, Ipswich, MA) pre-equilibrated in Buffer A, nutated for 1 hour at 4°C, then poured into a 2.5 cm diameter column. After the resin settled and the solution drained, it was washed with 100 ml wash buffer (Buffer A minus the Triton). Cleavage buffer (40 ml, wash buffer without Triton and with 50 mM DTT) was run through the resin at 4°C, then the column was plugged and moved to room temperature overnight. In the morning, 6 ml fractions were collected, eluting with 100 ml of wash buffer without Triton, and each was iced, assayed for protein, and the protein peak was pooled. The eluted Qc was aliquoted into PCR strips, snap frozen and stored at -80 °C. The N-terminal His_6_ tag was removed by TEV cleavage during the proteoliposome preparation process. Qc3Δ (Schwartz and Merz, 2009) was purified as above, except that the expression strain was Rosetta2(DE3), growth and IPTG induction were at 30°C, and Buffer A was replaced by Buffer B, composed of 20 mM TrisCl, pH 8.0, 0.5M NaCl, 10% glycerol and 1 mM EDTA. PMSF was added to 1 mM to the cell lysate. The chitin resin was nutated at 4°C for 3 hours, poured into the column, and the solution drained. The resin was washed with 30 ml of Buffer B, then 60 ml of Buffer B with 1.5M NaCl, then 60 ml of Buffer B, all at 4°C. Cleavage buffer was Buffer B + 50 mM DTT.

Plasmids expressing Sec17, Sec17LALA (Schwartz and Merz, 2009), Sec17KEKE and Sec17FSMS (Schwartz et al, 2017) were transformed into Rosetta2(DE3) and plated on LB+Ampicillin. A clone was grown overnight in 400 ml LB+Amp in a 2L flask with shaking at 37°C. In the morning, 125 ml portions were transferred into three 4 liter flasks which each had 2 liters of Terrific Broth with TB Salts and Amp. Cultures were grown to OD_600_ of 1.5, then shifted to an 18°C shaker where they received 0.5 mM IPTG and were shaken overnight. Cells were collected by centrifugation (JLA10.5 rotor, 5,000 rpm, 5 min, 3°C) and resuspended in 120 ml Buffer C (20 mM TrisCl, pH 8.0, 0.5 M KCl, 1 mM EDTA). PMSF (200 µM) and 0.01 volume of 100x PIC (100x protease inhibitor cocktail: 46 µg/ml leupeptin, 350 µg/ml pepstatin A, 240 µg/ml pefablock SC in dimethylsulfoxide) were added and the cells lysed by French press. The lysate was centrifuged (Beckman 60Ti rotor, 55,000 rpm, 30 min, 2°C). The supernatant was nutated with 60 ml of chitin resin previously equilibrated with Buffer C for 2 hours at 4°C, then poured into a 5 cm diameter column, drained, and washed with 500 ml Buffer C. Cleavage buffer (Buffer C + 40 mM DTT) was passed through the column (100 ml) and the column was plugged and moved to room temperature overnight. The next morning, 125 ml of Buffer C was added and 10 ml fractions were collected. The pooled protein fractions were concentrated in a YM-10 Centriprep, then dialyzed overnight into Rb150, aliquots frozen in liquid nitrogen, and stored at -80°C.

His_6_-Sec18 (Haas and Wickner 1996) was purified as follows. XL1 Blue/ pQE9-His_6_-Sec18 cells were grown in 300 ml LB + Ampicillin and Tetracycline at 37°C overnight with shaking. In the morning, three 90 ml portions were each added to 3 liters of LB with Amp and Tet in 6 L flasks and grown to an OD_600_ of 0.8. IPTG was added to 0.5 mM and growth continued for 4h to OD 1.25. Cells were collected (JLA10.5 rotor, 5,000 rpm, 5 min) and resuspended in 300 ml lysis buffer (100 mM HEPES-KOH, pH 7.0, 500 mM KCl, 5 mM ATP, 5 mM MgCl_2_, 0.014% ß-mercaptoethanol, 1x PIC) with 0.5 mM PMSF. Washed cells were sedimented a second time, then resuspended in 150 ml of the same lysis buffer with PMSF, then French pressed twice and centrifuged (Beckman 60Ti, 50,000 rpm, 1 h, 2°C). The cytosol was mixed with 12 ml of Qiagen Nickel-NTA resin in Buffer D (20 mM HEPES-KOH, pH 7.0, 480 mM KCl, 0.5 mM ATP, 10% glycerol, 1 mM MgCl_2_, 0.014% ß-mercaptoethanol, 20 mM ImidazoleCl), nutated for 1 hour at 4°C, then poured into a 1.5 cm column, drained, and washed with 100 ml Buffer 1. D. The protein was eluted with 50 ml of Buffer E (20 mM HEPES-KOH pH 7.0, 0.5 mM ATP, 10% glycerol, 1 mM MgCl_2_, 0.014% ß-mercaptoethanol, 500 mM ImidazoleCl) and 2 ml fractions were collected. The pooled protein was frozen in liquid nitrogen in 1 ml Eppendorf tubes and stored at -80°C for a later gel filtration step, as follows: Five 1 ml aliquots of the purified His_6_-Sec18 were thawed and centrifuged for 5 minutes at 4°C in a tabletop microfuge. They were pooled and applied to a 2.5 x 40 cm column with 196 ml of Superdex 200 prep grade resin (GE Healthcare, Uppsala, Sweden) in 20 mM PIPES-KOH, pH 6.8, 0.2 M sorbitol, 125 mM KCl, 5 mM MgCl_2_, 2 mM Na_2_ATP, 2 mM DTT, 10% glycerol at 4°C and 3ml fractions were collected. The gel filtered protein was pooled and snap frozen in PCR strips and stored at - 80°C.

DNA encoding the Qa, Qb, Qc and R SNARE Domains and transmembrane anchors, optimized for E. coli expression, was synthesized and cloned by Genewiz-Azenta (South Plainfield, NJ). All fragments were cloned into a pMBP-Parallel1 vector using the BamHI and SalI sites. The QaSNAREDomain-tm (QaSD-tm) includes both the 18 amino acids before the SNARE Domain of Vam3 which have been shown to be important for fusion (Song and Wickner, 2017) and its original transmembrane domain, amino acids 181-283. The construct QbSNAREDomain-tm corresponds to the amino acids 133-217 of Vti1 including the SNARE domain and its original transmembrane domain.

The construct QcSNAREDomain-tmQb includes the amino acids 259-316 of Vam7 followed by the transmembrane domain of Vti1, amino acids 195-217. The construct RSNAREDomain-tm includes the SNARE domain and original transmembrane domain of Nyv1, amino acids 165-253. For QcY42A-tm, DNA encoding the Qc SNARE containing the changed codon for amino acid 42 (Tyrosine to Alanine) and the transmembrane domain of the Qb SNARE was synthesized and cloned by Genewiz-Azenta. The fragment was codon optimized for expression in E. coli and cloned into the pMBP-Parallel1 vector using the BamHI and SalI sites. The final construct contains amino acids 1-316 of Vam7 followed by the transmembrane domain of Vti1, amino acids 195-217. All anchored SNARE domain proteins were expressed in BL21 STAR cells (Invitrogen, Waltham, MA) in the presence of 100μg/mL Ampicillin. The anchored QcY42A protein was expressed in Rosetta2(DE3)pLysS cells (Millipore Sigma, Burlington, MA) in the presence of 100μg/mL Ampicillin and 35μg/mL Chloramphenicol. The purification steps were identical for the anchored SNARE domains and QcY42A. Cells containing the MBP-tagged proteins were grown overnight in 100 ml of LB medium with antibiotics at 37ᵒC and shaking. In the morning, the 100 ml cell growth was poured into 3L of fresh medium plus antibiotics. Cells were grown to OD_600_ of 1, IPTG was added to 1 mM, and incubation with shaking continued for 5h. Cells were harvested by centrifugation (JLA10.5 rotor, 5,000 rpm, 5 min) and resuspended in 30 ml of 20 mM TrisCl, pH 8, 200 mM NaCl, 1 mM EDTA, 1 mM DTT, 0.5 mM PMSF and 1X PIC. Cell suspensions were French Pressed twice and centrifuged in a Beckman 60Ti rotor (50,000 rpm, 45 min, 4ᵒC). The membrane fraction, containing the anchored proteins, was resuspended in 23 ml of 20 mM TrisCl, pH 8, 200 mM NaCl, 1 mM EDTA, 1 mM DTT, 0.5 mM PMSF, 1X PIC and 1% Triton X100, nutated for 1h at 4ᵒC to solubilize the membranes and centrifugated again (50,000 rpm, 1h, 4°C). The supernatant was added to 15 ml of amylose resin equilibrated with 20 mM TrisCl, pH 8, 200 mM NaCl, 1 mM EDTA, 1 mM DTT and 1% Triton X100, nutated for 2h at 4ᵒC, and poured into a column. After the resin had settled and the column drained, the resin was washed with 100 ml of same buffer used for equilibration and protein was eluted with 40 ml of 20 mM HEPES-NaOH, pH 7.4, 200 mM NaCl, 10% glycerol, 1 mM DTT, 1% ß-octylglucoside and 10 mM maltose. The eluate was collected in 2 ml fractions and protein concentrations determined by Bradford. Small aliquots of proteins were frozen in liquid nitrogen and stored at -80ᵒC.

MBP-Qc-tm and the QcΔPX-tm variant (Xu and Wickner, 2012) were purified as described. HOPS and the soluble Vti1 (Zick and Wickner 2013), soluble Vam3 and Ypt7-tm (Song et al, 2020) and soluble GST-Nyv1 (Thorngren et al, 2004, Song et al, 2020) were purified as described.

Systematic names and ID numbers of all proteins used in this work according to Saccharomyces Genome Database (<u>Saccharomyces</u> <u>Genome Database | SGD</u>): Qa SNARE or Vam3 (YOR106W, SGD:S000005632), Qb SNARE or Vti1 (YMR197C, SGD:S000004810), Qc SNARE or Vam7 (YGL212W, SGD:S000003180), R SNARE or Nyv1 (YLR093C, SGD:S000004083), Rab or Ypt7 (YML001W, SGD:S000004460), Sec17 (YBL050W, SGD:S000000146), Sec18 (YBR080C, SGD:S000000284). HOPS subunits: Vps11 or Pep5 (YMR231W, SGD:S000004844), Vps16 or Vam9 (YPL045W, SGD:S000005966), Vps18 or Pep3 (YLR148W, SGD:S000004138), Vps33 or Pep14 (YLR396C, SGD:S000004388), Vps39 or Vam6 (YDL077C, SGD:S000002235), Vps41 or Vam2 (YDR080W, SGD:S000002487).

## Proteoliposome preparation

Proteoliposomes were prepared as in Zick and Wickner (2016) with modifications. For each 1 ml preparation of liposomes, 4 mM vacuolar mimic lipids in chloroform and 50 mM ß-octylglucoside were mixed in a glass vial. Vacuolar mimic lipid contains 47.6% 1,2-dilinoleoyl-*sn*-glycero-3-phosphocholine (Avanti 18:2 PC item 850385C), 18% 1,2-dilinoleoyl-*sn*-glycero-3-phosphoethanolamine (Avanti 18:2 PE item 850755C), 18% L-α-phosphatidylinositol (Avanti soy PI item 840044C), 4.4% 1,2-dilinoleoyl-*sn*-glycero-3-phospho-L-serine (Avanti 18:2 PS item 840040C), 2% 1,2-dilinoleoyl-*sn*-glycero-3-phosphate (Avanti 18:2 PA, item 840885C), 8% ergosterol (Fluka Sigma item 45480-10G-F), 1% 1,2-dipalmitoyl-*sn*-glycerol (Avanti 16:0 DG item 800816C), and 1% Di-*myo*-phosphatidylinositol 3-phosphate (Echelon PI(3)P, diC16, item P-3016, dissolved in 1:2:0.8 chloroform:methanol:water). Lipid detergent mixtures were dried under a stream of nitrogen for 30 min. Solvent was further removed by centrifugation under vacuum for 3 hours. Lipid pellets were resuspended by nutation at room temperature in 400 μl of 50 mM HEPES-NaOH, pH 7.4, 375 mM NaCl, 25% glycerol, 2.5 mM MgCl_2_.

To prepare proteoliposomes for fusion assays, SNAREs were added to the 400 μl of reconstituted lipids at a 1:16,000 SNARE to lipid molar ratio, and the Rab, when added, was at a 1:8,000 molar ratio to lipid. His_6_-TEV (1 µM) was added to each preparation to remove purification tags. Suspensions with the R SNARE also received 250 μl of 16 µM R-phycoerythrin solution (Invitrogen item P811), while suspensions with any Q SNARE received 250 μl of 32 µM Cy5-streptavidin solution (SeraCare item 5270-0023). Water was added to bring the volume of each preparation to 1 ml, but was mixed with the lipid-detergent solution before proteins or fluorescent probes were added. To prepare liposomes for assays of ATP hydrolysis or lipid binding, the fluorescent probes were omitted, their volumes replaced by water, and the SNAREs, when present, were added at a 1:2,000 protein to lipid ratio. For lipid binding assays, 1% of the 18:2 PC was replaced by 1% Rhodamine-DHPE (Invitrogen item L1392). The lipid-detergent-protein solutions were nutated for 30 min, 4°C, then dialyzed overnight, in the dark, at 4°C in 12 mm (flat width) 25kD MWCO dialysis tubing (Repligen Item 132550), against 200 ml (per liposome prep) of Rb150+Mg (20 mM HEPES-NaOH, pH 7.4, 150 mM NaCl, 10% glycerol, 1 mM MgCl_2_) with 1 g BioBeads SM-2 (BioRad 152-3920) per 200 ml dialysis buffer. Liposomes were harvested into 2 ml Eppendorf tubes on ice, mixed with an equal volume of 70% Histodenz in modified Rb150+Mg (containing only 2% glycerol) and transferred to Beckman 11 x 60 mm ultraclear centrifuge tubes. These were overlaid with 25% Histodenz in modified Rb150+Mg to the 3.8 ml mark on the tube. A final layer of 600 µl Rb150+Mg was added to the top. Tubes were centrifuged (60Ti, 55,000 rpm, 90 min, 4°C). Liposomes were harvested onto ice and lipid phosphorus concentration determined (Chen, et al., 1956). Liposomes were diluted with Rb150+Mg to 2 mM lipid phosphate, aliquoted (30 μl) into PCR strips, and frozen and stored in liquid nitrogen.

## Binding assays

Assays of proteins bound to liposomes were conducted as described (Orr and Wickner, 2022). Liposomes were prepared with VML lipids and 1% Rhodamine-DHPE as described above. In brief, 30 μl reactions in Rb150 contained 0.5mM lipid, 0.04% defatted BSA, 1 mM MgCl_2_, and soluble proteins as indicated. These were incubated in PCR strips in a 27° C water bath for 1 hour. Reactions received 90 μl of 54% Histodenz in modified Rb150+Mg with only 2% glycerol and were gently vortexed. A portion (80 μl) of each was transferred to 7 x 20 mm ultracentrifuge tubes, over-layered with 80 μl of 35%, then 80 μl of 30% Histodenz in modified Rb150+Mg, then 50 µl of Rb150+Mg. The gradients were centrifuged in a Beckman TLS55 swinging bucket rotor (55,000 rpm, 30 min, 4°C). The top 80 μl was harvested, solubilized by adding Thesit to 0.125%, nutated for 30 minutes, and the percentage of recovery determined by Rhodamine fluorescence, as compared to the likewise solubilized starting samples. Samples were boiled in SDS-PAGE sample buffer with ß-mercaptoethanol and analyzed by Western blot. Assays were performed in triplicate and UN-SCAN-IT software (Silk Scientific, Orem, Utah) was used to quantify bands. Averages and standard deviations are presented.

## ATP hydrolysis assays

ATP hydrolysis was assayed with a phosphate assay kit ab270004 from Abcam (Cambridge, UK) according to the manufacturer’s instructions. Before the assay, Sec18 was freed from ATP and phosphate with an Amicon Ultra 50K 0.5mL Centrifugal Filter (Merck-Millipore, Burlington, MA) as follows: His_6_-Sec18 (100μl) was diluted with 400 μl buffer (20 mM PIPES-KOH, pH6.8, 200 mM sorbitol, 125 mM KCl, 5 mM MgCl_2_, 2 mM DTT and 10% glycerol) and centrifuged (10,000 x g, 6 min, 4ᵒC) to return to approximately the original volume of 100 μl. This was repeated five times, using a total of 2 ml of this buffer. To collect the final phosphate and ATP-free His_6_-Sec18, the filter was inserted upside down in a clean collection tube and centrifugated (1000 x g, 2 min, 4ᵒC). Protein concentration was determined by Bradford reagent.

ATP hydrolysis reactions (20 μl) were in 0.5 ml conical plastic tubes for 10-30 min at 22°C, terminated by the addition of 20 μl of 6 mM EDTA. Proprietary solutions were added as per manufacturer’s instructions. Reactions were transferred to wells of a clear, 384 well, square well microplate (item 781101, Greiner Bio-One, Kremsmunster, Austria) and read in a BioTek EPOCH plate reader (Winooski, VT) at Ab650. All assays were performed in triplicate. Averages and standard deviations are shown, with relevant statistics (T-test analysis).

## Acknowledgements

This work was supported by NIH grant 2R35GM118037. We thank Gus Lienhard, Jeremy Thorner, Alex Merz and Axel Brunger for insightful comments.

## Supplemental Material

The Rab Ypt7 and the lipid PI3P are required for fusion of vacuoles (Haas et al., 1995; Seeley et al., 2002) or of vacuole-derived proteoliposomes (Stroupe et al., 2009; Mima and Wickner, 2009). Do they directly affect the ATPase activity of Sec18? Liposomes were prepared with or without PI3P and with or without Rab, and these four proteoliposome preparations were examined in systematic assays of ATP hydrolysis with either single components present (Sec18, Sec17, Qc-SNARE, or proteoliposomes with or without Rab and with or without PI3P), with pairs of Sec18 and one additional component, with combinations of three, or with all four (Sec18, Sec17, Qc, and proteoliposomes) present (Supplemental Figure S1). Modest concentrations of each Sec18 stimulant were employed to keep the highest stimulated levels of ATP hydrolysis within the assay range. Individual components (Supplemental Figure S1, lanes 1, 2, 3, 7, 8) including Sec18 alone (lane 3) gave little signal. Addition of any of the four kinds of proteoliposomes to Sec18 gave comparable small stimulations (lanes 9, 10, 19, 20), suggesting that neither PI3P nor Ypt7 stimulated Sec18 directly. Qc gave little further stimulation in any case (lanes 11, 12, 21, 22). There was also no difference between these four proteoliposomes in stimulating ATPase activity with Sec17 (lanes 13, 14, 23, 24; no Qc). PI3P (but not Ypt7) was needed for maximal stimulation of Sec18 ATPase in the presence of Sec17, proteoliposomes, and Qc (lanes 15, 16, 25, 26). Since only HOPS and Qc (among the vacuole fusion factors) have direct affinity for PI3P (Stroupe et al., 2006; Cheever et al., 2001), and HOPS is not needed to activate the Sec18 ATPase (Figure 3; Figure 6B below), the sole function of PI3P for activating Sec18 ATPase is as a Qc receptor.

**Supplemental Figure S1.**
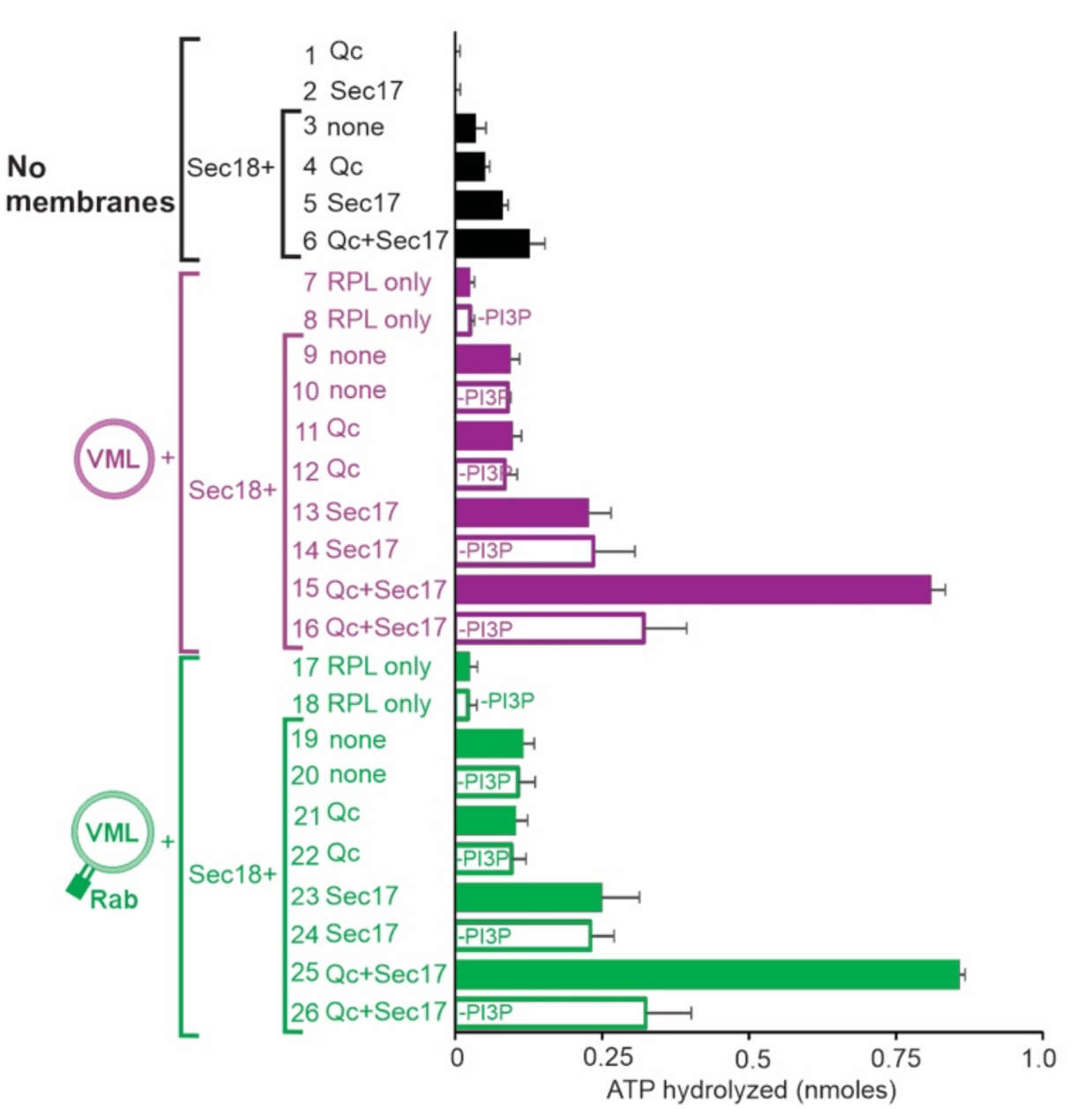
Together, Sec17, Qc, and liposomes stimulate Sec18 more than any subsets. PI3P is only needed for Sec18 ATP hydrolysis in the presence of Qc, Sec17, and liposomes.

**Supplemental Figure S2.**
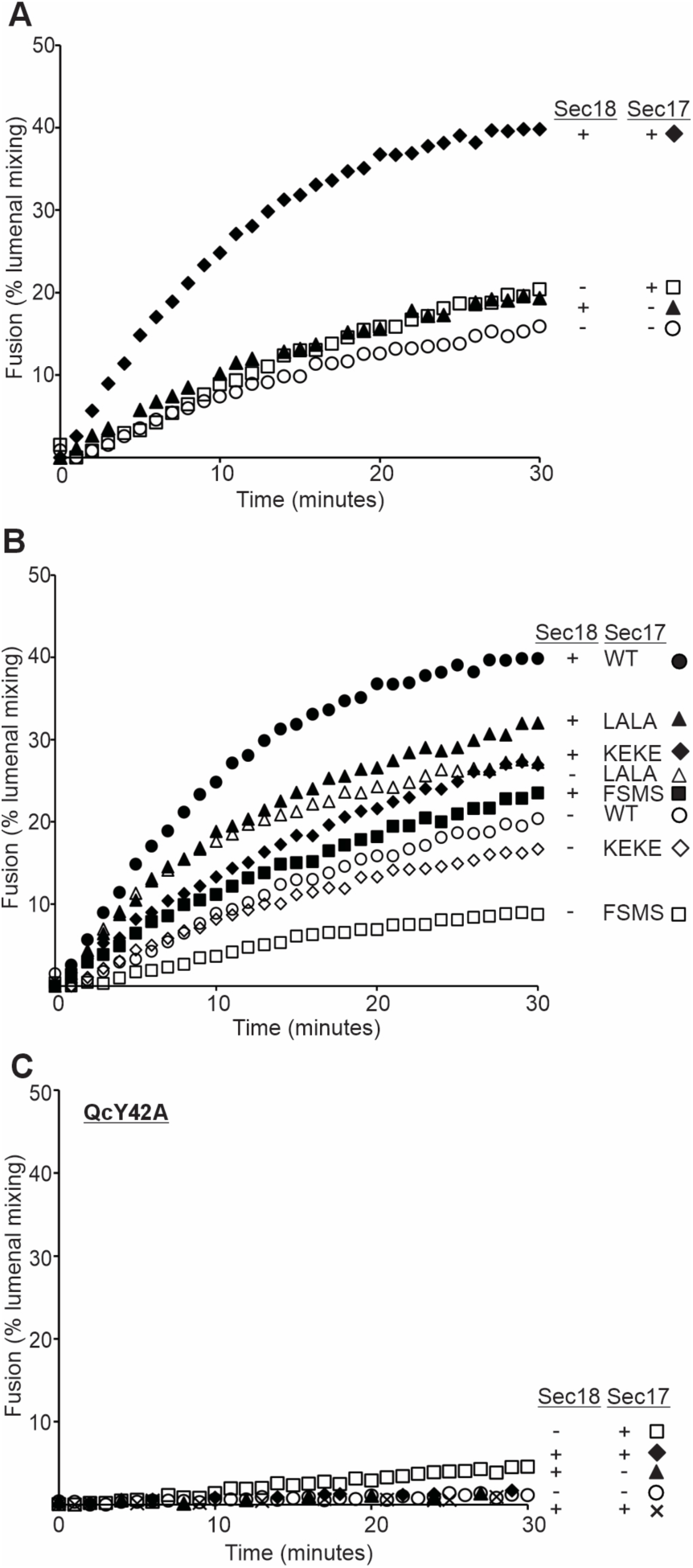
(A) and (B) Kinetics are shown for a representative set of assays from Figure 6A. (C) Qc recognition of PI3P is needed for fusion. Fusion assays were as in A, but with 20 nM QcY42A instead of wild-type Qc. The symbol x indicates omission of Qc. Assays were performed with or without 100 nM wild-type Sec17 and with or without 50 nM Sec18.

